# Humoral immunity induced by XEC monovalent vaccines against SARS-CoV-2 variants including XEC, LP.8.1, NB.1.8.1, XFG, and BA.3.2

**DOI:** 10.64898/2025.12.11.693810

**Authors:** Keiya Uriu, Yu Kaku, Yusuke Kosugi, Luo Chen, Naoya Itoh, Yoshifumi Uwamino, Hiroshi Fujiwara, Hironori Satoh, The Genotype to Phenotype Japan (G2P-Japan) Consortium, Kei Sato

**Affiliations:** Division of Systems Virology, Department of Microbiology and Immunology, The Institute of Medical Science, The University of Tokyo, Tokyo, Japan; Department of Clinical Infectious Diseases, Graduate School of Medical Sciences, Nagoya City University, Aichi, Japan; Nagoya City University East Medical Center, Aichi, Japan; Graduate School of Medicine, The University of Tokyo, Tokyo, Japan; Graduate School of Frontier Sciences, The University of Tokyo, Chiba, Japan; Department of Infectious Diseases, Graduate School of Medical Sciences, Nagoya City University, Aichi, Japan; Department of Laboratory Medicine, Keio University School of Medicine, Tokyo, Japan; Fujiwara Clinic, Tokyo, Japan; Satoh Clinic, Tokyo, Japan; International Vaccine Design Center, The Institute of Medical Science, The University of Tokyo, Tokyo, Japan; Collaboration Unit for Infection, Joint Research Center for Human Retrovirus infection, Kumamoto University, Kumamoto, Japan; MRC-University of Glasgow Centre for Virus Research, Glasgow, UK; Faculty of Medicine, Chulalongkorn University, Bangkok, Thailand; Programme in Emerging Infectious Diseases, Duke-NUS Medical School, Singapore

## Abstract

After the emergence of SARS-CoV-2 Omicron variant at the end of 2021, Omicron has highly diverged into various sublineages: e.g., BA.5 in 2022, XBB.1.5 in 2023, JN.1 in 2024. Currently, several JN.1 subvariants including XEC, LP.8.1, NB.1.8.1 and XFG are circulating worldwide. Additionally, recent studies show that BA.3.2, a descendant of Omicron BA.3, exhibits profound immune evasion potential. On December 5, 2025, BA.3.2, was designated a variant under monitoring by the WHO.

To prevent COVID-19, several countries including Japan have continuously developed and approved variant-adapted vaccines: e.g., ancestral/BA.5 bivalent vaccine in 20223, XBB.1.5-based monovalent vaccine in 20234, JN.1-based monovalent vaccine in 2024, and LP.8.1-based monovalent vaccine in 2025. Since XEC was more prevalent than LP.8.1 at the beginning of 2025 in Japan, two Japanese pharmaceutical companies, Daiichi Sankyo and Meiji Seika Pharma, have produced XEC-based vaccines. Here, we investigated the antiviral immunity induced by XEC-based monovalent vaccines against recently circulating SARS-CoV-2 variants, as well as BA.3.2, in the Japanese population.

To assess the neutralizing antibody response induced by XEC-based vaccines, we obtained sera from individuals who had been vaccinated with the XEC-based mRNA monovalent vaccine produced by Daiichi Sankyo (N=22) or the XEC-based self-amplifying replicon vaccine produced by Meiji Seika Pharma (N=20). We collected sera before and 3–4 weeks after vaccination and then performed a neutralization assay using these sera with lentivirus-based pseudoviruses harboring spike proteins of B.1.1, BA.5, XBB.1.5, JN.1, XEC, LP.8.1, NB.1.8.1, XFG and BA.3.2. The 50% neutralization titers of the sera against all variants tested were significantly increased in after vaccination in both Daiichi Sankyo cohort (2.1-fold to 11.9-fold, P<0.0001) and Meiji Seika Pharma cohort (1.4-fold to 3.9-fold, P<0.0005). Consistent with our recent study using the sera from LP.8.1-based vaccinees, the humoral immunity induced by XEC-based vaccines against JN.1 and its subvariants (XEC, LP.8.1, NB.1.8.1 and XFG) was greater than that against other variants (B.1.1, BA.5, XBB.1.5 and BA.3.2). Notably, our result showed that the XEC-based mRNA vaccine induces a stronger humoral immunity compared to the XEC-based replicon vaccine, and even the LP.8.1-adapted vaccines. However, it should be noted that our cohorts are relatively small and may include confounding factors that affect the results, such as age, sex, history of natural infection and vaccination status. Future investigations with larger cohorts are required to better understand this possibility.

To compare the immune status induced by XEC- and LP.8.1-based vaccines, we analyzed the cross-neutralization induced by these vaccines using antigenic cartography. The antigenic map was depicted based on the 50% neutralization titers values obtained from this study and our recent study using LP.8.1-based vaccine sera. The cartography showed that the immune status induced by XEC-based vaccines was similar to that induced by LP.8.1-based vaccines against the nine SARS-CoV-2 variant antigens tested. In sum, our investigations, including the recent one8, suggest that all 4 of the JN.1 subvariants-based vaccines that we tested induce profound humoral immunity against a broad range of SARS-CoV-2 variants.

## Text

After the emergence of SARS-CoV-2 Omicron variant at the end of 2021, Omicron has highly diverged into various sublineages: e.g., BA.5 in 2022, XBB.1.5 in 2023, JN.1 in 2024. Currently, several JN.1 subvariants including XEC, LP.8.1, NB.1.8.1 and XFG are circulating worldwide.^1^ Additionally, recent studies show that BA.3.2, a descendant of Omicron BA.3, exhibits profound immune evasion potential.^2^ On December 5, 2025, BA.3.2, was designated a variant under monitoring by the WHO (https://www.who.int/activities/tracking-SARS-CoV-2-variants).

To prevent COVID-19, several countries including Japan have continuously developed and approved variant-adapted vaccines: e.g., ancestral/BA.5 bivalent vaccine in 2022^3^, XBB.1.5-based monovalent vaccine in 2023^4^, JN.1-based monovalent vaccine in 2024^5,6^ and LP.8.1-based monovalent vaccine in 2025.^7,8^ Since XEC was more prevalent than LP.8.1 at the beginning of 2025 in Japan, two Japanese pharmaceutical companies, Daiichi Sankyo and Meiji Seika Pharma, have produced XEC-based vaccines. Here, we investigated the antiviral immunity induced by XEC-based monovalent vaccines against recently circulating SARS-CoV-2 variants, as well as BA.3.2, in the Japanese population.

To assess the neutralizing antibody response induced by XEC-based vaccines, we obtained sera from individuals who had been vaccinated with the XEC-based mRNA monovalent vaccine produced by Daiichi Sankyo (N=22; **Figure A**) or the XEC-based self-amplifying replicon vaccine produced by Meiji Seika Pharma (N=20; **Figure B**). We collected sera before and 3–4 weeks after vaccination and then performed a neutralization assay using these sera with lentivirus-based pseudoviruses harboring spike proteins of B.1.1, BA.5, XBB.1.5, JN.1, XEC, LP.8.1, NB.1.8.1, XFG and BA.3.2. The 50% neutralization titers of the sera against all variants tested were significantly increased in after vaccination in both Daiichi Sankyo cohort (2.1-fold to 11.9-fold, *P*<0.0001) (**Figure A**) and Meiji Seika Pharma cohort (1.4-fold to 3.9-fold, *P*<0.0005) (**Figure B**). Consistent with our recent study using the sera from LP.8.1-based vaccinees^8^, the humoral immunity induced by XEC-based vaccines against JN.1 and its subvariants (XEC, LP.8.1, NB.1.8.1 and XFG) was greater than that against other variants (B.1.1, BA.5, XBB.1.5 and BA.3.2) (**Figure**). Notably, our result showed that the XEC-based mRNA vaccine (**Figure A**) induces a stronger humoral immunity compared to the XEC-based replicon vaccine (**Figure B**), and even the LP.8.1-adapted vaccines.^8^ However, it should be noted that our cohorts are relatively small and may include confounding factors that affect the results, such as age, sex, history of natural infection and vaccination status. Future investigations with larger cohorts are required to better understand this possibility.

**Figure.**
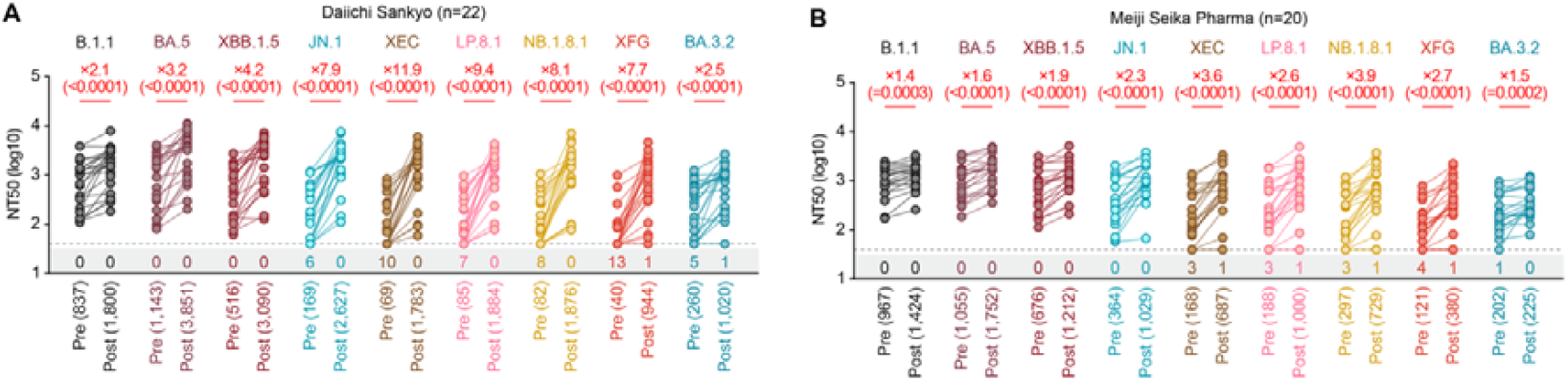
The cross-neutralization induced by XEC monovalent vaccine. Neutralization assay. Assays were performed with pseudoviruses harboring the spike proteins of SARS-CoV-2 B.1.1, BA.5, XBB.1.5, JN.1, XEC, LP.8.1, NB.1.8.1, XFG and BA.3.2. The following two sera were used: XEC monovalent vaccine sera from fully vaccinated individuals who had received Daiichi Sankyo XEC vaccine (Daiichi Sankyo cohort) (one 3-dose vaccinated, one 4-dose vaccinated, one 5-dose vaccinated, two 6-dose vaccinated, nine 7-dose vaccinated and eight 8-dose vaccinated; twenty two donors in total, average age: 65.6, range: 28–84, 27.3% male) (**A**), and those from fully vaccinated individuals who received Meiji Seika Pharma XEC vaccine (Meiji Seika Pharma cohort) (three 5-dose vaccinated, eight 6-dose vaccinated, four 7-dose vaccinated, three 8-dose vaccinated and two 9-dose vaccinated; twenty donors, average age: 44.4, range: 24–62, 45.0% male) (**B**). Assays for each serum sample were performed in quadruplicate to determine the 50% neutralization titer (NT50). Each dot represents one NT50 value for each donor, and the NT50 values for the same donor before (“Pre”) and 3–4 weeks after (“Post”) vaccination are connected by a line. Numbers in parentheses below the graphs indicate the median of the NT50 value. The horizontal dashed lines indicate the lowest serum dilution of 40-fold. The number of sera with the NT50 values below the lower detection limit is shown in the figure (under the bottom horizontal dashed line). The limit of detection was calculated as 1.5 shown as a gray area. The fold changes between Pre and Post indicated in red with “X” are the median of paired NT50s obtained from each individual. The p-values in the parentheses were determined by two-sided Wilcoxon signed-rank tests (**A and B**).

To compare the immune status induced by XEC- and LP.8.1-based vaccines, we analyzed the cross-neutralization induced by these vaccines using antigenic cartography. The antigenic map was depicted based on the 50% neutralization titers values obtained from this study and our recent study using LP.8.1-based vaccine sera.^8^ The cartography showed that the immune status induced by XEC-based vaccines was similar to that induced by LP.8.1-based vaccines against the nine SARS-CoV-2 variant antigens tested (**Supplemental figure**). In sum, our investigations, including the recent one^8^, suggest that all 4 of the JN.1 subvariants-based vaccines that we tested induce profound humoral immunity against a broad range of SARS-CoV-2 variants.

## Grants

Supported in part by KAKENHI (25K10373, to Yu Kaku); Nagoya Co-Creation Research Fund (202502006, to Yu Kaku); the Outstanding Research Group Support Program in Nagoya City University (2530003, to Naoya Itoh); the Department of Clinical Infectious Diseases, Nagoya City University Graduate School of Medical Sciences (to Naoya Itoh); AMED ASPIRE program (25jf0126002 to Kei Sato); AMED SCARDA Japan Initiative for World-leading Vaccine Research and Development Centers “UTOPIA” (JP223fa627001 to Kei Sato); AMED SCARDA Program on R&D of new generation vaccine including new modality application (253fa727002 to Kei Sato); AMED Research Program on Emerging and Re-emerging Infectious Diseases (24fk0108907, 25fk0108690 to Kei Sato); AMED Japan Program for Infectious Diseases Research and Infrastructure (Collaborative Research via Overseas Research Centers) (25wm0225041 to Kei Sato); AMED Japan Program for Infectious Diseases Research and Infrastructure (Collaborative Research via Overseas Research Centers) (25wm0225041 to Kei Sato) ; JSPS KAKENHI fund for the Promotion of Joint International Research (International Leading Research) (JP23K20041 to Kei Sato); JSPS KAKENHI grant-in-aid for Scientific Research A (JP24H00607 to Kei Sato); Mitsubishi UFJ Financial Group, Inc. Vaccine Development grant to Kei Sato); Takeda Science Foundation( to Kei Sato) and JP24ama121012 (S02820001 and S02820002 to Kei Sato).

