## Supplementary Appendix for "Humoral immunity induced by XEC monovalent vaccines against SARS-CoV-2 variants including XEC, LP.8.1, NB.1.8.1, XFG, and BA.3.2"

**Table of Contents**

**Contents** Page


| | |
|---|---|
| **Materials and Methods** | 2-3 |
| Ethics statement | |
| Human serum collection | |
| Cell culture | |
| Pseudovirus preparation | |
| Neutralization assay | |
| **Table S1.** Human vaccination sera used in this study | 4 |
| **Table S2.** Summary of cohort information | 5 |
| **Table S3.** Raw data of 50% neutralization titer | 6 |
| **Supplemental figure** | 7 |
| **Consortia** | 8–9 |
| **Acknowledgments** | 10 |
| **Supplemental References** | 11 |

### Materials and Methods

#### Ethics statement

All protocols involving specimens from human subjects recruited at Keio University, Fujiwara Clinic, and Satoh Clinic were reviewed and approved by the Institutional Review Boards of The Institute of Medical Science, The University of Tokyo (approval IDs: 2021-1-0416 and 2022-29-0915), and Keio University (approval ID: 20200059), respectively. All human subjects provided written informed consent. All protocols for the use of human specimens were reviewed and approved by the Institutional Review Boards of The Institute of Medical Science, The University of Tokyo (approval IDs: 2021-1-0416, 2021-18-0617 and 2022-29-0915).

#### Human serum collection

XEC monovalent vaccine sera from fully vaccinated individuals who had received Daiichi-Sankyo XEC vaccine (Daiichi Sankyo cohort) (one 3-dose vaccinated, one 4-dose vaccinated, one 5-dose vaccinated, two 6-dose vaccinated, nine 7-dose vaccinated and eight 8-dose vaccinated; twenty two donors in total, average age: 65.6, range: 28–84, 27.3% male) and those from fully vaccinated individuals who received Meiji Seika Pharma XEC vaccine (Meiji Seika Pharma cohort) (three 5-dose vaccinated, eight 6-dose vaccinated, four 7-dose vaccinated, three 8-dose vaccinated and two 9-dose vaccinated; twenty donors, average age: 44.4, range: 24–62, 45.0% male). For this study, we collected samples before vaccination and three to four weeks (21–28 days) after vaccination.
Sera were inactivated at 56°C for 30 minutes and stored at –80°C until use. The details of the vaccine sera are summarized in **Table S1 and S2**.

#### Cell culture

The Lenti-X 293T cells (Takara, Cat# 632180) and HOS-ACE2/TMPRSS2 cells (kindly provided by Dr. Kenzo Tokunaga), a derivative of HOS cells (a human osteosarcoma cell line; ATCC CRL-1543) stably expressing human ACE2 and TMPRSS2[1,2] were maintained in Dulbecco’s modified Eagle's medium (DMEM) (high glucose) (Wako, Cat# 044- 29765) containing 10% fetal bovine serum (Sigma-Aldrich Cat# 172012-500ML), 100 units penicillin and 100 ug/ml streptomycin (Sigma-Aldrich, Cat# P4333-100ML).

#### Pseudovirus preparation

Plasmids expressing the SARS-CoV-2 spike (S) proteins of B.1.1, BA.5, XBB.1.5, JN.1, XEC, LP.8.1, NB.1.8.1, XFG and BA.3.2 were prepared in our previous studies.[3-10] Pseudoviruses were prepared as previously described.[6-10] Briefly, lentivirus (HIV-1)-based, luciferase-expressing reporter viruses were pseudotyped with the SARS-CoV-2 S.

One prior day of transfection, the LentiX-293T cells were seeded at a density of $2 \times 10^6$ cells. The LentiX-293T cells were cotransfected with 1 μg psPAX2-IN/HiBiT (a packaging plasmid encoding the HiBiT-tag-fused integrase[1], 1 μg pWPI-Luc2 (a reporter plasmid encoding a firefly luciferase gene[11] and 500 ng plasmids expressing parental S or its derivatives using TransIT-293 transfection reagent (Mirus, Cat# MIR2704) according to the manufacturer's protocol. Two days post transfection, the culture supernatants were harvested and filtrated. The pseudoviruses were harvested and stored at –80°C until use after filtration.

**Neutralization assay**

Neutralization assays were performed previously described[6-10] and mainly conducted by a semi-automated high-throughput method using Fluent780 (Tecan).[6-10,12] The SARS-CoV-2 spike pseudoviruses (counting ~100,000 relative light units) and serially diluted (40-fold to 29,160-fold dilution at the final concentration) heat-inactivated sera were manually prepared in a 2-ml 96-well plate (Greiner, Cat# 780271) and in 96-well microplates (ThermoFisher Scientific, Cat# 168136), respectively. The pseudoviruses were dispensed and mixed with the sera in 384-well plates (ThermoFisher Scientific, Cat# 164610) on Fluent780 (Tecan). Pseudoviruses without sera were included as controls. After incubation at 37°C for 1 hour, HOS-ACE2/TMPRSS2 cells (6,000 cells/30 μl) were added to the 20 μl mixture of pseudovirus and serum in the 384-well white plate on the device. Two days post infection, the infected cells were lysed with a Bright-Glo luciferase assay system (Promega, Cat# E2620) on Fluent780 (Tecan), and the luminescent signal was measured and processed using an Infinite200 and a Magellan (Tecan). The assay of each serum sample was performed in quadruplicate, and the 50% neutralization titer ($NT_{50}$) was calculated using Prism 9 (GraphPad Software). The limit of detection (LOD) is determined to be 30 based on our previous report.[10] The raw data of $NT_{50}$ in this study were summarized in **Table S3**.

**Antigenic cartography**

Based on the 50% neutralization titers, the antigenic cartography (**Supplementary Figure**) of the viruses and vaccine sera was depicted with the method described by Smith *et al.*[13] using the Racmacs v1.1.35 on R v4.2.2 (https://www.r-project.org/) with 500 optimization runs.

**Table S1. Human vaccination sera used in this study**

| XEC vaccine manufacturers | Donor ID | Sex | Age | Date of 1st vaccination (YYYY-MM-DD) | Date of 2nd vaccination (YYYY-MM-DD) | Date of 3rd vaccination (YYYY-MM-DD) | Date of 4th vaccination (YYYY-MM-DD) | Date of 5th vaccination (YYYY-MM-DD) | Date of 6th vaccination (YYYY-MM-DD) | Date of 7th vaccination (YYYY-MM-DD) | Date of 8th vaccination (YYYY-MM-DD) | Date of 9th vaccination (YYYY-MM-DD) | Date of sampling (before vaccination) (YYYY-MM-DD) | Date of XEC vaccination (YYYY-MM-DD) | Date of sampling (after vaccination) (YYYY-MM-DD) | Time interval between vaccination and the second sampling | Prior infection? (YYYY-MM-DD) |
|---|---|---|---|---|---|---|---|---|---|---|---|---|---|---|---|---|---|
| Daiichi-Sankyo | F000100 | Male | 45 | 2021-04-28 (P) | 2021-05-20 (P) | 2022-01-08 (P) | 2022-07-15 (P) | 2022-12-03 (P) | 2023-10-27 (P) | 2024-10-01 (D,JN1) | - | - | 2025-10-07 | 2025-10-07 | 2025-10-29 | 22 | No |
| Daiichi-Sankyo | F000200 | Female | 46 | 2021-04-28 (P) | 2021-05-19 | 2021-12-25 (P) | 2022-06-15 (P) | 2023-10-25 (P) | 2024-10-03 (D,JN1) | - | - | - | 2025-10-02 | 2025-10-02 | 2025-10-23 | 21 | No |
| Daiichi-Sankyo | F000300 | Male | 63 | 2021-07-03 (M) | 2021-07-31 (M) | 2022-03-12 (P) | 2022-08-18 (M) | 2022-11-21 (P) | 2024-10-04 (D) | - | - | - | 2025-10-01 | 2025-10-01 | 2025-10-22 | 21 | Yes (2023-07) |
| Daiichi-Sankyo | F000400 | Female | 82 | 2021-05-20 (P) | 2021-06-10 (P) | 2022-02-10 (P) | NA | NA | NA | NA | 2024-11-11 (P) | - | 2025-10-04 | 2025-10-04 | 2025-10-27 | 23 | No |
| Daiichi-Sankyo | F000500 | Female | 54 | 2021-05-20 (P) | 2021-06-10 (P) | 2022-01-28 (P) | 2022-08-05 (M) | 2022-12-14 (P) | 2023-06-21 (M) | 2023-11-14 (P) | 2024-11-11 (P) | - | 2025-10-04 | 2025-10-04 | 2025-10-27 | 23 | No |
| Daiichi-Sankyo | F000868 | Female | 75 | 2021-06 (P) | 2021-07 (P) | 2022-02-08 (P) | 2022-07-15 (P) | 2022-11-18 (P) | 2023-06-06 (P) | 2023-10-13 (P) | - | - | 2025-10-03 | 2025-10-03 | 2025-10-27 | 24 | No |
| Daiichi-Sankyo | F018678 | Female | 63 | NA | NA | 2022-02-15 (P) | 2022-07-15 (P) | 2022-11-01 (P) | 2023-06-27 (P) | 2023-09-29 (P) | 2024-10-02 (D,JN1) | - | 2025-10-03 | 2025-10-03 | 2025-10-25 | 22 | Yes (2023-03, 2024-08) |
| Daiichi-Sankyo | F023012 | Female | 82 | 2021-05-21 (P) | 2021-06-08 (P) | 2022-01-28 (P) | 2022-07-01 (P) | 2022-11-11 (P) | 2023-05-26 (P) | 2023-11-07 (P) | - | - | 2025-10-07 | 2025-10-07 | 2025-10-28 | 21 | No |
| Daiichi-Sankyo | F035951 | Female | 74 | NA | NA | 2022-02-18 (P) | 2022-07-19 (P) | 2023-01-30 (P) | 2023-06-13 (P) | 2024-10-04 (D,JN1) | - | - | 2025-10-03 | 2025-10-03 | 2025-10-24 | 21 | No |
| Daiichi-Sankyo | F037838 | Female | 70 | 2021-06-29 (P) | 2021-07-25 (P) | 2022-02-19 (P) | 2022-08-09 (P) | 2022-12-13 (P) | 2023-07-11 (P) | 2023-11-14 (P) | 2024-10-01 (D,JN1) | - | 2025-10-01 | 2025-10-01 | 2025-10-23 | 22 | No |
| Daiichi-Sankyo | F039982 | Female | 67 | 2021-06-25 (P) | 2021-07-16 (P) | 2022-02-18 (P) | 2022-07-19 (P) | 2022-12-05 (P) | 2023-10-13 (P) | 2024-11-12 (P) | - | - | 2025-10-01 | 2025-10-01 | 2025-10-22 | 21 | No |
| Daiichi-Sankyo | F050041 | Male | 74 | 2021-06-15 (P) | 2021-07-16 (P) | 2022-02-18 (P) | 2022-07-19 (P) | 2022-11-18 (P,BA4/5) | 2023-06-13 (P,BA4/5) | 2024-12-06 (P) | - | - | 2025-10-07 | 2025-10-07 | 2025-10-28 | 21 | No |
| Daiichi-Sankyo | F050351 | Female | 79 | NA | NA | NA | 2022-07-19 (P) | 2022-11-08 (P) | 2023-05-16 (P) | 2023-09-26 (P) | 2024-10-04 (D,JN1) | - | 2025-10-07 | 2025-10-07 | 2025-10-29 | 22 | No |
| Daiichi-Sankyo | F052203 | Male | 80 | 2021-06-13 (M) | 2021-07-14 (M) | 2022-01-28 (P) | 2022-11-01 (P) | 2023-05-23 (P) | 2023-10-17 (P) | 2024-11-05 (P) | - | - | 2025-10-03 | 2025-10-03 | 2025-10-28 | 25 | No |
| Daiichi-Sankyo | F055336 | Female | 75 | 2021-06-11 (P) | 2021-07-02 (P) | 2022-02-04 (P) | 2022-07-08 (P) | 2022-11-15 (P) | 2023-05-19 (P) | 2023-10-06 (P) | 2024-10-02 (D,JN1) | - | 2025-10-01 | 2025-10-01 | 2025-10-24 | 21 | No |
| Daiichi-Sankyo | F072023 | Female | 57 | 2021-07-05 (P) | 2021-07-26 (P) | 2022-02-10 (M) | 2022-07-15 (P) | 2022-12-23 (P,BA4/5) | 2023-07-14 (P,BA1) | 2023-11-10 (XBB) | 2024-10-01 (D,JN1) | - | 2025-10-03 | 2025-10-03 | 2025-10-24 | 21 | Yes (2024-02 2025-02) |
| Daiichi-Sankyo | F072088 | Female | 71 | 2021-06-24 (P) | 2021-07-15 (P) | 2022-03-04 (P) | 2022-08-25 (P) | 2023-01-06 (P) | 2023-05-16 (P) | 2023-09-26 (P) | 2024-10-03 (D,JN1) | - | 2025-10-04 | 2025-10-04 | 2025-10-29 | 25 | No |
| Daiichi-Sankyo | F075080 | Female | 75 | 2021-06-08 (P) | 2021-06-29 (P) | 2022-07-12 (P) | 2022-11-08 (P) | 2023-05-16 (P) | 2023-09-29 (P) | 2024-10-01 (D,JN1) | - | - | 2025-10-01 | 2025-10-01 | 2025-10-22 | 21 | No |
| Daiichi-Sankyo | F078713 | Female | 28 | 2021-04 (P) | 2021-04 (P) | 2021-12 (P) | - | - | - | - | - | - | 2025-10-03 | 2025-10-03 | 2025-10-24 | 21 | Yes (2022-11) |
| Daiichi-Sankyo | F079661 | Male | 62 | NA | NA | NA | 2022-10 (P) | 2024-10-01 (D,JN1) | - | - | - | - | 2025-10-03 | 2025-10-03 | 2025-10-24 | 21 | No |
| Daiichi-Sankyo | F079996 | Male | 84 | NA | NA | NA | NA | 2023-06-13 (P) | 2023-10-24 (P) | 2024-10-04 (D,JN1) | - | - | 2025-10-03 | 2025-10-03 | 2025-10-24 | 21 | Yes (2023-01) |
| Daiichi-Sankyo | F082525 | Female | 37 | NA | NA | NA | 2022-10 (P) | - | - | - | - | - | 2025-10-03 | 2025-10-03 | 2025-10-28 | 25 | Yes (2022-08) |
| Meiji Seika Pharma | S0015 | Female | 24 | 2021-06-18 (P) | 2021-07-09 (P) | 2022-03-31 (P) | 2022-09-29 (P) | 2023-10-08 (P) | - | - | - | - | 2025-10-04 | 2025-10-04 | 2025-10-27 | 23 | Yes (2022-01, 2023-07) |
| Meiji Seika Pharma | S0019 | Male | 37 | 2021-06-19 (M) | 2021-07-17 (M) | 2022-05-13 (M) | 2022-10-28 (M) | 2023-10-06 (M) | 2024-05-11 (M) | 2024-08-03 (P) | 2024-10-20 (MSP) | - | 2025-09-28 | 2025-09-28 | 2025-10-24 | 26 | No |
| Meiji Seika Pharma | S0049 | Female | 34 | 2021-01-01 (P) | NA | NA | NA | NA | NA | NA | NA | 2025-10-20 (MSP) | 2025-09-29 | 2025-09-29 | 2025-10-20 | 21 | Yes (2023-11-01) |
| Meiji Seika Pharma | S0066 | Male | 41 | 2021-07-15 (M) | 2021-08-12 (M) | 2022-02-15 (M) | 2022-10-14 (P) | 2023-10-16 (P) | 2024-11-04 (MSP) | - | - | - | 2025-10-04 | 2025-10-04 | 2025-10-25 | 21 | Yes (2023-07-29, 2025-07-18) |
| Meiji Seika Pharma | S0074 | Female | 49 | 2021-07-10 (P) | 2021-07-30 (P) | 2022-02-10 (M) | 2022-10-06 (M) | 2023-09-29 (P) | 2024-11-16 (MSP) | - | - | - | 2025-09-27 | 2025-09-27 | 2025-10-18 | 21 | No |
| Meiji Seika Pharma | S0087 | Female | 45 | 2021-08-21 (M) | 2021-09-18 (M) | 2022-03-30 (P) | 2024-03-01 (D) | 2024-12-20 (MSP) | - | - | - | - | 2025-09-26 | 2025-09-26 | 2025-10-20 | 24 | Yes (2022-08-04, 2025-07-04) |
| Meiji Seika Pharma | S0094 | Female | 49 | 2021-05-01 (P) | 2021-05-22 (P) | 2022-02-06 (P) | 2022-08-04 (P) | 2023-05-11 (P) | 2023-10-26 (P) | 2024-11-01 (MSP) | - | - | 2025-09-29 | 2025-09-29 | 2025-10-27 | 28 | Yes (2022-12-06) |
| Meiji Seika Pharma | S0112 | Male | 59 | 2021-08-16 (P) | 2021-09-06 (P) | 2022-03-07 (P) | 2022-10-25 (P) | 2023-10-10 (P) | 2024-06-21 (M) | 2024-10-07 (M) | 2025-04-08 (P) | - | 2025-09-26 | 2025-09-26 | 2025-10-17 | 21 | Yes (2024-02, 2024-09-04) |
| Meiji Seika Pharma | S0119 | Male | 36 | 2021-07-19 (M) | 2021-08-23 (M) | 2022-03-09 (M) | 2023-11-10 (P) | 2023-10-27 (M) | 2024-11-22 (MSP) | - | - | - | 2025-10-01 | 2025-10-01 | 2025-10-27 | 26 | Yes (2023-09-04, 2025-08-02) |
| Meiji Seika Pharma | S0135 | Male | 40 | 2021-07-02 (P) | 2021-07-23 (P) | 2022-02-16 (M) | 2022-11-02 (P) | 2023-09-22 (P) | 2024-11-03 (MSP) | - | - | - | 2025-09-26 | 2025-09-26 | 2025-10-21 | 25 | No |
| Meiji Seika Pharma | S0140 | Female | 48 | 2021-07-18 (M) | 2021-08-15 (M) | 2022-02-26 (M) | 2022-10-24 (M) | 2023-11-24 (P) | 2024-11-12 (MSP) | - | - | - | 2025-09-29 | 2025-09-29 | 2025-10-27 | 28 | No |
| Meiji Seika Pharma | S0162 | Female | 62 | 2021-07-16 (P) | 2021-08-06 (P) | 2022-02-28 (M) | 2022-10-04 (M) | 2023-10-20 (M) | 2024-10-21 (P) | - | - | - | 2025-09-29 | 2025-09-30 | 2025-10-28 | 28 | Yes (2023-04-02) |
| Meiji Seika Pharma | S0171 | Male | 40 | 2021-07-08 (M) | 2021-08-03 (M) | 2022-04-21 (P) | 2022-12-02 (M) | 2023-10-16 (P) | 2024-10-24 (P) | - | - | - | 2025-09-28 | 2025-09-28 | 2025-10-25 | 27 | Yes (2022-02-18) |
| Meiji Seika Pharma | S0185 | Female | 51 | 2021-06-24 (M) | 2021-07-23 (M) | 2022-02-15 (M) | 2022-07-15 (P) | 2022-11-03 (P) | 2023-06-02 (P) | 2023-11-07 (P) | 2024-07-19 (M) | 2024-11-01 (MSP) | 2025-09-30 | 2025-09-30 | 2025-10-21 | 21 | No |
| Meiji Seika Pharma | S0194 | Male | 27 | 2021-08-29 (P) | 2021-09-18 (P) | 2022-03-22 (M) | 2022-11-11 (P) | 2023-09-23 (P) | 2024-11-16 (P) | 2025-04-12 (MSP) | - | - | 2025-09-28 | 2025-09-28 | 2025-10-22 | 24 | No |
| Meiji Seika Pharma | S0197 | Female | 41 | 2021-07-04 (P) | 2021-07-25 (P) | 2022-01-31 (M) | 2022-10-02 (M) | 2023-09-29 (M) | 2024-05-31 (P) | 2024-10-03 (P) | 2025-03-19 (MSP) | - | 2025-09-27 | 2025-09-27 | 2025-10-20 | 23 | No |
| Meiji Seika Pharma | S0219 | Female | 57 | 2021-07-12 (M) | 2021-08-09 (M) | 2022-02-15 (M) | 2022-10-08 (P) | 2023-10-08 (M) | - | - | - | - | 2025-09-29 | 2025-09-29 | 2025-10-20 | 21 | No |
| Meiji Seika Pharma | S0221 | Male | 45 | 2021-08-01 (P) | 2021-08-22 (P) | 2022-03-06 (M) | 2022-10-10 (M) | 2023-10-28 (M) | 2024-06-14 (M) | 2024-10-20 (MSP) | - | - | 2025-09-28 | 2025-09-28 | 2025-10-25 | 27 | Yes (2023-08-30) |
| Meiji Seika Pharma | S0235 | Female | 49 | 2021-08-25 (P) | 2021-09-24 (P) | 2022-04-07 (N) | 2022-12-01 (P) | 2023-11-22 (P) | 2024-11-16 (MSP) | - | - | - | 2025-09-29 | 2025-09-29 | 2025-10-21 | 22 | No |
| Meiji Seika Pharma | S0261 | Male | 54 | 2021-07-14 (P) | 2021-08-12 (P) | 2022-02-24 (M) | 2022-10-28 (P) | 2023-06-15 (P) | 2023-12-28 (P) | 2024-11-15 (MSP) | - | - | 2025-09-26 | 2025-09-26 | 2025-10-20 | 24 | No |

NA, not applicable.

A, Astrazeneca; P, Pfizer/BioNTech; M, Moderna; S, Sinovac; J, Jenner; N, Novavax/Takeda; D, Daiichi-Sankyo; MSP, Meji Seika Pharma

BA1/2, BA.1/2 bivalent vaccine; BA4/5, BA.4/5 bivalent vaccine; XBB, XBB.1.5 monovalent vaccine; JN1, JN.1 monovalent vaccine

**Table S2. Raw data of 50% neutralization titer**

| XEC-based vaccine | | B.1.1 | | BA.5 | | XBB.1.5 | | JN.1 | | XEC | | LP.8.1 | | NB.1.8.1 | | XFG | | BA.3.2 | |
|---|---|---|---|---|---|---|---|---|---|---|---|---|---|---|---|---|---|---|---|
| manufacturers | Donor ID | Pre | Post | Pre | Post | Pre | Post | Pre | Post | Pre | Post | Pre | Post | Pre | Post | Pre | Post | Pre | Post |
| Daiichi-Sankyo | F000100 | 615.1 | 3055 | 551.6 | 5566 | 463.5 | 4607 | 156.1 | 2963 | 81.15 | 3209 | 91.24 | 3380 | 40 | 3010 | 40 | 1853 | 105 | 1195 |
| Daiichi-Sankyo | F000200 | 1959 | 3003 | 2931 | 5960 | 1320 | 4789 | 673.7 | 2697 | 464.3 | 2450 | 521.2 | 2252 | 412.3 | 2093 | 259.9 | 1412 | 669.9 | 1255 |
| Daiichi-Sankyo | F000300 | 2251 | 2221 | 3712 | 4556 | 2717 | 3550 | 1152 | 2556 | 601.1 | 1487 | 564.5 | 1371 | 556.8 | 1565 | 110.6 | 1036 | 989.3 | 1064 |
| Daiichi-Sankyo | F000400 | 116.8 | 184.6 | 80.38 | 200.5 | 136.6 | 126.4 | 40 | 144.7 | 40 | 96.35 | 40 | 102.3 | 40 | 99.64 | 40 | 107.2 | 40 | 40 |
| Daiichi-Sankyo | F000500 | 3867 | 3766 | 917.4 | 1218 | 127.5 | 426.4 | 40 | 2369 | 40 | 1597 | 40 | 1538 | 40 | 1836 | 40 | 1874 | 830.3 | 737.1 |
| Daiichi-Sankyo | F000868 | 657.5 | 1789 | 1368 | 4868 | 934.5 | 4526 | 346.4 | 2843 | 40 | 825.8 | 79.73 | 869.5 | 65.63 | 646.4 | 40 | 292.9 | 285.6 | 1148 |
| Daiichi-Sankyo | F018678 | 1066 | 1136 | 2385 | 3655 | 1841 | 2433 | 1118 | 2182 | 840.2 | 2124 | 962.2 | 2098 | 1052 | 2194 | 997 | 2297 | 614.5 | 901 |
| Daiichi-Sankyo | F023012 | 112.5 | 331.1 | 98.7 | 608.9 | 108.8 | 477.8 | 40 | 305.9 | 40 | 90.78 | 40 | 77.37 | 40 | 88.64 | 40 | 40 | 40 | 208.3 |
| Daiichi-Sankyo | F035951 | 1017 | 1481 | 1411 | 3959 | 567.8 | 2868 | 477.4 | 3125 | 266.1 | 2551 | 294.9 | 2185 | 263.7 | 2013 | 84.83 | 346.3 | 446.7 | 1176 |
| Daiichi-Sankyo | F037838 | 1019 | 3377 | 1679 | 11237 | 923 | 6023 | 316.2 | 7233 | 40 | 4515 | 63.04 | 3963 | 74.83 | 5126 | 40 | 2607 | 377.9 | 2613 |
| Daiichi-Sankyo | F039982 | 109.3 | 683.6 | 84.11 | 989.3 | 62 | 729 | 40 | 1074 | 40 | 847.1 | 40 | 765.9 | 40 | 903.2 | 40 | 314.6 | 40 | 224.2 |
| Daiichi-Sankyo | F050041 | 1266 | 3612 | 1432 | 8617 | 801.2 | 7305 | 132.1 | 3879 | 69.46 | 3082 | 99.78 | 3295 | 81.74 | 2751 | 40 | 1622 | 233.5 | 1891 |
| Daiichi-Sankyo | F050351 | 1256 | 1810 | 2428 | 3743 | 1859 | 3312 | 1178 | 7683 | 274.6 | 5953 | 369.9 | 4152 | 287.1 | 7101 | 86.32 | 3242 | 564.1 | 1008 |
| Daiichi-Sankyo | F052203 | 355.6 | 962.7 | 302.6 | 1736 | 155.8 | 1429 | 51.74 | 1033 | 40 | 167.4 | 40 | 236.9 | 40 | 99.02 | 40 | 61.24 | 49.67 | 351.6 |
| Daiichi-Sankyo | F055336 | 378 | 415.1 | 193 | 294.7 | 72.74 | 147.2 | 40 | 110.8 | 40 | 58.66 | 40 | 77.42 | 40 | 76.7 | 40 | 53.52 | 136.9 | 112.4 |
| Daiichi-Sankyo | F072023 | 132.4 | 282.2 | 125.6 | 716.8 | 135.2 | 680 | 182.2 | 1291 | 106.5 | 996.4 | 99.15 | 776.2 | 114 | 752.4 | 94.93 | 440 | 40 | 141.5 |
| Daiichi-Sankyo | F072088 | 3867 | 7878 | 4132 | 10557 | 1642 | 6290 | 452.9 | 4367 | 185.3 | 4051 | 196.8 | 3088 | 152.3 | 3110 | 96.82 | 1215 | 1236 | 2672 |
| Daiichi-Sankyo | F075080 | 212.9 | 323.1 | 405 | 869.1 | 366.4 | 823.2 | 105.9 | 893.9 | 40 | 571.8 | 68.69 | 549 | 86.41 | 714.3 | 86.83 | 649.7 | 74.39 | 173.5 |
| Daiichi-Sankyo | F078713 | 176.6 | 3116 | 80.49 | 7233 | 104.9 | 4320 | 40 | 2825 | 40 | 1542 | 40 | 1955 | 40 | 1412 | 40 | 523.1 | 40 | 1871 |
| Daiichi-Sankyo | F079661 | 1324 | 1517 | 2064 | 3219 | 1572 | 2580 | 1132 | 2340 | 684.7 | 1968 | 645.6 | 1812 | 688.4 | 1916 | 561.3 | 1396 | 428 | 501.5 |
| Daiichi-Sankyo | F079996 | 1367 | 2868 | 1931 | 7390 | 992.5 | 3971 | 580.9 | 3321 | 255.9 | 2435 | 210.1 | 2326 | 302.7 | 2420 | 40 | 852.9 | 480.8 | 1032 |
| Daiichi-Sankyo | F082525 | 321 | 3228 | 335.3 | 8720 | 262.4 | 5768 | 104.5 | 3829 | 69.19 | 4316 | 64.84 | 4329 | 82.05 | 4476 | 40 | 4648 | 108 | 1582 |
| Meiji Seika Pharma | S0015 | 392.5 | 660.5 | 402.1 | 917.9 | 189.2 | 650.4 | 129.6 | 657.4 | 99.7 | 427.1 | 109.3 | 749 | 99.49 | 455.1 | 40 | 282.3 | 154.1 | 257.2 |
| Meiji Seika Pharma | S0019 | 780.4 | 869.6 | 450.8 | 556.1 | 113.4 | 210.5 | 57.81 | 67.17 | 40 | 40 | 40 | 40 | 40 | 40 | 40 | 40 | 64.4 | 80.81 |
| Meiji Seika Pharma | S0049 | 977.3 | 1630 | 1774 | 4021 | 2090 | 3232 | 1436 | 3812 | 1158 | 3332 | 1881 | 4932 | 909.5 | 3696 | 781.9 | 2041 | 578.2 | 1247 |
| Meiji Seika Pharma | S0066 | 1929 | 2729 | 3215 | 3851 | 3151 | 3156 | 1145 | 2280 | 521.3 | 959.5 | 610.1 | 1167 | 771.4 | 1372 | 225.9 | 447 | 581.8 | 750.8 |
| Meiji Seika Pharma | S0074 | 1323 | 1823 | 1240 | 2296 | 696.5 | 1489 | 401.4 | 1976 | 194.4 | 1298 | 287.5 | 1894 | 297.6 | 1834 | 207.2 | 1441 | 521.9 | 921.8 |
| Meiji Seika Pharma | S0087 | 1616 | 1958 | 3224 | 4959 | 2302 | 2523 | 2082 | 2947 | 1380 | 2736 | 1771 | 2362 | 1190 | 2548 | 415.3 | 862.7 | 974.5 | 1158 |
| Meiji Seika Pharma | S0094 | 905.1 | 796.8 | 709.2 | 1099 | 360.5 | 752.7 | 254 | 877.1 | 116.8 | 618.4 | 148.6 | 1027 | 105.1 | 791.4 | 59.41 | 500.7 | 148.8 | 309.6 |
| Meiji Seika Pharma | S0112 | 870.4 | 1112 | 870.5 | 1641 | 477.1 | 1016 | 381.8 | 2202 | 260.8 | 2247 | 244.9 | 2763 | 295.7 | 2372 | 133 | 2276 | 190 | 345.3 |
| Meiji Seika Pharma | S0119 | 2489 | 1947 | 2674 | 2274 | 1599 | 1661 | 1236 | 1171 | 563.7 | 635.8 | 678.1 | 1021 | 524.2 | 616.7 | 170.6 | 198.3 | 792.2 | 672.2 |
| Meiji Seika Pharma | S0135 | 1688 | 1357 | 1599 | 1719 | 934.8 | 1110 | 449.8 | 872.5 | 136.3 | 672.9 | 153 | 320.5 | 132.2 | 784 | 89.58 | 349.8 | 506.6 | 440.6 |
| Meiji Seika Pharma | S0140 | 1821 | 2105 | 521.5 | 601.2 | 166.9 | 303.4 | 746.6 | 1072 | 832.9 | 991.8 | 1005 | 1702 | 573.9 | 683.4 | 588.9 | 712.9 | 134 | 132.6 |
| Meiji Seika Pharma | S0162 | 182 | 563.2 | 701.1 | 2746 | 655.3 | 2111 | 157.9 | 943.9 | 91.88 | 425.6 | 120.5 | 496.3 | 83.47 | 309.9 | 117.8 | 321.6 | 40 | 392.6 |
| Meiji Seika Pharma | S0171 | 1275 | 1624 | 1341 | 1785 | 1063 | 1649 | 293.3 | 501.2 | 142.5 | 428.1 | 195.5 | 482.4 | 297.5 | 453.4 | 124.3 | 226.8 | 317.8 | 325.6 |
| Meiji Seika Pharma | S0185 | 771.1 | 1491 | 599 | 1602 | 315.3 | 1129 | 179.6 | 979.1 | 78.25 | 700.4 | 96.23 | 415.2 | 93.45 | 759.6 | 83.9 | 329.2 | 233.6 | 456.6 |
| Meiji Seika Pharma | S0194 | 957.3 | 1239 | 1313 | 1852 | 965.7 | 1294 | 1248 | 2217 | 563.5 | 1291 | 803.7 | 2041 | 557.6 | 1409 | 478.5 | 1174 | 213.2 | 185.9 |
| Meiji Seika Pharma | S0197 | 172.9 | 260.7 | 186.1 | 360.6 | 164 | 307.6 | 346.8 | 1205 | 266.9 | 1283 | 179.9 | 1015 | 315.4 | 1412 | 169.9 | 741.9 | 105.7 | 248.5 |
| Meiji Seika Pharma | S0219 | 566.2 | 1117 | 300.9 | 620 | 210 | 910.9 | 63.98 | 527.4 | 40 | 248.5 | 40 | 282.2 | 40 | 246 | 40 | 81.44 | 115.3 | 248.7 |
| Meiji Seika Pharma | S0221 | 2101 | 3288 | 3443 | 4551 | 2230 | 5230 | 1100 | 2212 | 486.2 | 1195 | 635.8 | 698.7 | 328.2 | 698 | 114.1 | 321.3 | 586.8 | 774.1 |
| Meiji Seika Pharma | S0235 | 1280 | 1841 | 1535 | 2532 | 795.6 | 1442 | 257.9 | 342 | 40 | 70.03 | 40 | 86.63 | 40 | 77.32 | 40 | 40.03 | 184.7 | 213.2 |
| Meiji Seika Pharma | S0261 | 567.9 | 779.6 | 310 | 895.6 | 216.9 | 644.8 | 72.23 | 985.1 | 83.37 | 540.1 | 92.31 | 985.8 | 63.8 | 564.6 | 82.11 | 410.1 | 85.38 | 227.8 |

**Table S3. Cohort comparison summary**

| Vaccine strain | Vaccine manufacturers | Average age | % male | Average times of vaccination | Average times of infection |
|---|---|---|---|---|---|
| XEC | Daiichi Sankyo | 65.6 | 27.3 | 6.9 | 0.4 |
| XEC | Meiji Seika Pharma | 44.4 | 45.0 | 6.7 | 0.8 |
| LP.8.1 | Pfizer/BioNTech | 33.8 | 37.9 | 5.2 | 0.8 |
| LP.8.1 | Novavax | 44.5 | 45.0 | 6.4 | 0.5 |

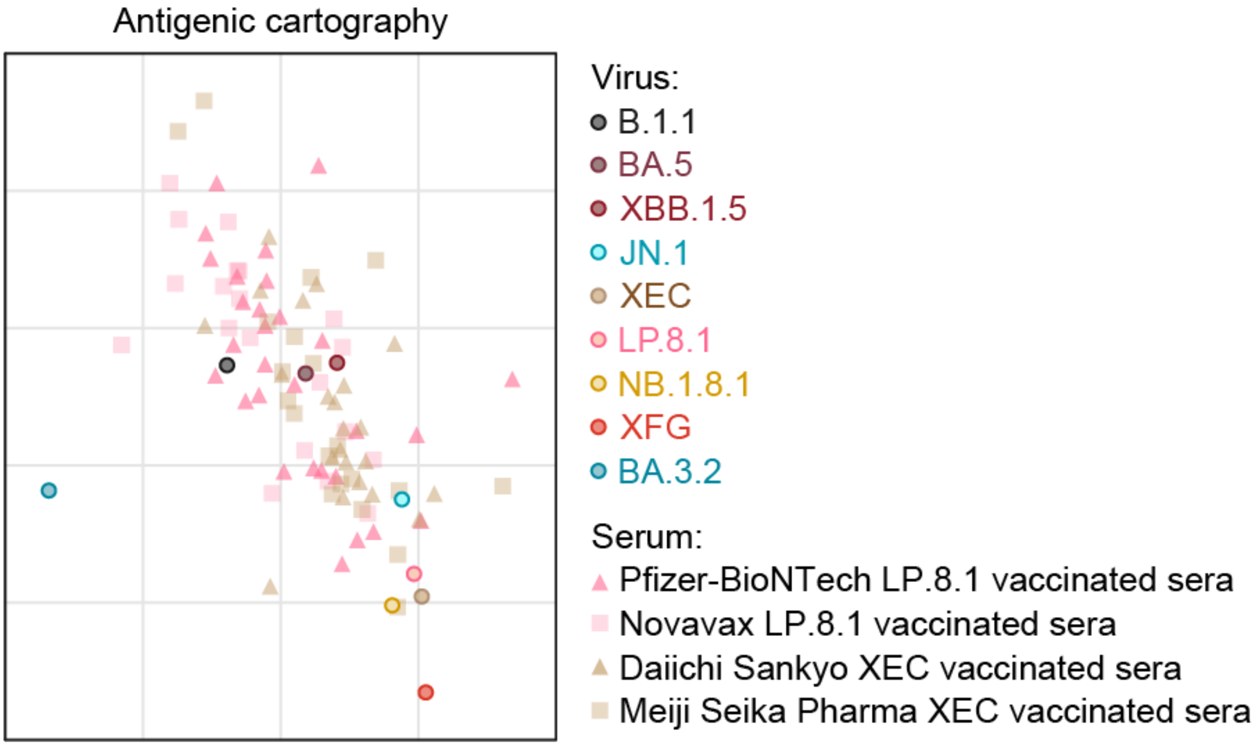


**Supplemental figure. Antigenic cartography**
Antigenic map based on the serum neutralization data from the neutralization assay. Virus positions are represented by closed circles. Both axes represent antigenic distance with one antigenic distance unit (AU) in any direction corresponding to a 2-fold change in NT50.

### Consortia

#### The Genotype to Phenotype Japan (G2P-Japan) Consortium

**The Institute of Medical Science, The University of Tokyo, Japan**
Jumpei Ito, Naoko Misawa, Arnon Plianchaisuk, Ziyi Guo, Kaoru Usui, Wilaiporn Saikruang, Spyridon Lytras, Shusuke Kawakubo, Luca Nishimura, Mohammad Golam Kibbria, Shigeru Fujita, Jarel Elgin M. Tolentino, Yukun Zhu, Ananporn Supataragul, Moonjong Kang, Wenye Li, Maximilian Stanley Yo, Yueying Zhang, Mizuka Fujiwara, Mai Suganami, Mika Chiba, Shiho Tanaka, Eiko Ogawa, Kaho Okumura, Tsuki Fukuda, Tamaki Yoshihara, Keiko Koizumi, Mizuho Ota, Hiroaki Unno, Hinata Yokokawa, Yuka Ozaki, Nanako Kanai, Kyoko Kurihara

**Hokkaido University, Japan**
Takasuke Fukuhara, Tomokazu Tamura, Rigel Suzuki, Saori Suzuki, Shuhei Tsujino, Hayato Ito, Hirofumi Sawa, Naganori Nao, Keita Matsuno, Keita Mizuma, Jingshu Li, Izumi Kida, Yume Mimura, Yuma Ohari, Shinya Tanaka, Masumi Tsuda, Lei Wang, Yoshikata Oda, Zannatul Ferdous, Kenji Shishido, Hiromi Mohri, Miki Iida

**Tokyo Metropolitan Institute of Public Health**
Kenji Sadamasu, Kazuhisa Yoshimura, Hiroyuki Asakura, Isao Yoshida, Mami Nagashima

**Tokai University, Japan**
So Nakagawa

**Kyoto University, Japan**
Kotaro Shirakawa, Akifumi Takaori-Kondo, Kazuo Takayama, Rina Hashimoto, Sayaka Deguchi, Yukio Watanabe, Yoshitaka Nakata, Hiroki Futatsusako, Ayaka Sakamoto, Naoko Yasuhara, Takao Hashiguchi, Tateki Suzuki, Kanako Kimura, Jiei Sasaki, Yukari Nakajima, Hisano Yajima

**Hiroshima University, Japan**
Takashi Irie, Ryoko Kawabata

**Kyushu University, Japan**
Kaori Tabata

**Kumamoto University, Japan**

Terumasa Ikeda, Hesham Nasser, Ryo Shimizu, MST Monira Begum, Michael Jonathan, Yuka Mugita, Sharee Leong, Otowa Takahashi, Takamasa Ueno, Chihiro Motozono, Mako Toyoda

**University of Miyazaki, Japan**
Akatsuki Saito, Anon Kosaka, Miki Kawano, Natsumi Matsubara, Tomoko Nishiuchi

**Charles University, Czechia**
Jiri Zahradnik, Prokopios, Andrikopoulos, Miguel Padilla-Blanco, Aditi Konar, Ruojin Tuan

**Acknowledgments**

We would like to thank all members of The Genotype to Phenotype Japan (G2P-Japan) Consortium. We thank Kenzo Tokunaga (National Institute of Infectious Diseases, Japan) for sharing materials, Yuka Kamoshita (Department of Laboratory Medicine, Keio University School of Medicine, Japan), Masayo Noguchi (Clinical Laboratory, Keio University Hospital, Japan) for supporting patient sera collection, and Mika Chiba, Tsuki Fukuda, Mizuho Ota, Tamaki Yoshihara, Keiko Koizumi (Division of Systems Virology, University of Tokyo, Japan) for performing experimental assays.
